# Split-ends is modulating lipid droplet content in adult *Drosophila* glial cells and is protective against paraquat toxicity

**DOI:** 10.1101/2020.05.20.101220

**Authors:** Victor Girard, Valérie Goubard, Matthieu Querenet, Laurent Seugnet, Laurent Pays, Serge Nataf, Eloïse Dufourd, David Cluet, Bertrand Mollereau, Nathalie Davoust

**Affiliations:** Laboratory of Biology and Modelling of the Cell, UMR5239 CNRS, ENS de Lyon, UMS 344 Biosciences Lyon Gerland, Université de Lyon, France; CarMeN Laboratory, INSERM UMR-1060, INRA U1235, Université Claude Bernard Lyon1, INSA of Lyon, Charles Merieux Medical School, Université de Lyon, France; Banque de Tissus et de Cellules des Hospices Civils de Lyon, Hôpital Edouard Herriot, Lyon, France; Centre de Recherche en Neurosciences de Lyon, Equipe physiologie intégrée du système d’éveil, UMR5292, INSERM U1028, Université Claude Bernard Lyon1, France; Institut Universitaire de France

## Abstract

Glial cells are early sensors of neuronal injury and are able to store lipids in lipid droplets under oxidative stress conditions. Here, we investigated the glial functions of Spen in the context of Parkinson’s disease (PD). Using a data mining approach, we first found that the human ortholog of *spen*, *SPEN/SHARP* belongs to the set of astrocyte-expressed genes which mRNA levels are significantly different in the *substantia nigra* of PD patients as compared to controls. Interestingly, the retrieved list of differentially expressed genes was enriched in genes involved in lipid metabolism. In a *Drosophila* model of PD, we observed that *spen* mutant flies were more sensitive to paraquat intoxication. Moreover, the glia-restricted knockdown of *spen* led to the expansion and the accumulation of lipid droplets as well as the inhibition of Notch pathway. Taken together our results show that Spen regulates lipid metabolism and storage in glial cells and by these means contribute to glia-mediated functions in the context of neurodegeneration.

## INTRODUCTION

Parkinson’s disease (PD) is a neurodegenerative disorder characterized by the selective loss of dopaminergic neurons in the *substantia nigra pars compacta*. Specific environmental factors combined to a permissive genetic background are thought to trigger PD. Among environmental clues, chronic exposure to pesticide has been demonstrated to be involved in the aetiology of PD. While genetic models of PD are available in *Drosophila*, the paraquat-induced model reproduces several important pathophysiological features that currently provide causal links between chronic pesticide intoxication and PD. In particular, oxidative stress and ROS production in the brain of paraquat-intoxicated flies were shown to induce the degeneration of dopaminergic neurons, leading to severe motor disability and precocious death^1–3^. In a large range of species, from *Drosophila* to humans, glial cells are early sensors of central nervous system (CNS) injury and respond to neuronal damages by morphological modifications, proliferation and the engagement of specific transcriptomic programs^4^. We and others recently demonstrated that the hallmark of glial response to stress includes the intra-cytoplasmic accumulation of lipid droplets^5–9^.

Interestingly, Spen is a glia-expressed protein, which, in other cell types, has been implicated in lipid metabolism and the formation of lipid droplets^10^. During *Drosophila* development, Spen has been shown to regulate midline glia specification in the embryo^11^ and survival of inter-ommatidial glial cells in pupal retina^12^. Spen proteins are characterized by two functional domains that are conserved along evolution: an RNA binding motifs (RRM, RNA Recognition Motif) and a Spen Paralog and Ortholog C-terminal (SPOC) domain^11,13–15^. Spen mediates its biological effects through transcriptional regulation, RNA silencing and splicing or via epigenetic mechanisms relying on the direct interaction of Spen with chromatin^16–21^.

Of note also, Oswald et al. have described the human homologue of *spen*, *SPEN/SHARP* as a novel component of the Notch/RBP-Jkappa signalling pathway^22^. Moreover, a recent structural study demonstrated that RBP-J can indeed bind to SPEN and act as co-repressor^23^. Accordingly, in *Drosophila,* Spen antagonises Notch signalling during pupal retinal cells development^24^. However, it has also been shown that SPEN can recruit the KMT2D co-activator complex to Notch target genes^25^ and a study suggests that Spen could act upstream of Notch receptor activation by regulating the trafficking of the Notch ligand Delta in intestinal stem cells of adult flies.

On this basis, we investigated the role of Spen in adult *Drosophila* glia as a putative regulator of lipid metabolism and Notch signalling pathway, and under paraquat exposure as a model of PD.

## RESULTS

### *SPEN*, the human ortholog of *spen*, is up-regulated in the *substantia nigra pars compacta* of PD patients

We first sought to determine whether *SPEN* (also named *SHARP),* the human ortholog of *spen*, had been reported to be expressed by human astrocytes and if its expression was differentially regulated in the context of PD. To this aim, we took advantage of a recent meta-analysis of microarray datasets obtained from *substantia nigra* samples derived from PD vs control subjects^26^. From the list of differentially expressed genes (data supplement 1), we then performed a tissue enrichment analysis using the TargetMine webtool^27^. We found that *SPEN* belongs to a set of 197 genes up-regulated in the SN of PD patients and previously shown to be expressed by human astrocytes^28^ (data supplement 2). On another hand, 469 astrocyte-expressed genes were identified as being down-regulated in the SN of PD patients (data supplement 2). It should be noticed however that such an observation is not a formal proof that *SPEN* is up-regulated in the astrocytes of PD patients. Indeed, *SPEN* is also expressed by neurons in the human brain^28^. Interestingly, we also found that the whole list of genes identified as being constantly down-regulated in the *substantia nigra* of PD patients^26^ was significantly enriched in lipid-related genes (data supplement 3 and table 1) according to the NIH (National Institute of Health)-run pathway enrichment webtool BioPlanet^29^. No specific enrichment was observed when assessing the whole list of up-regulated genes (data supplement 4). Altogether, these data mining results support, at least in part, the implication of *SPEN* in PD and lead us to investigate the effect of *spen* expression in the *Drosophila* paraquat model of PD described below.

**Table 1.**
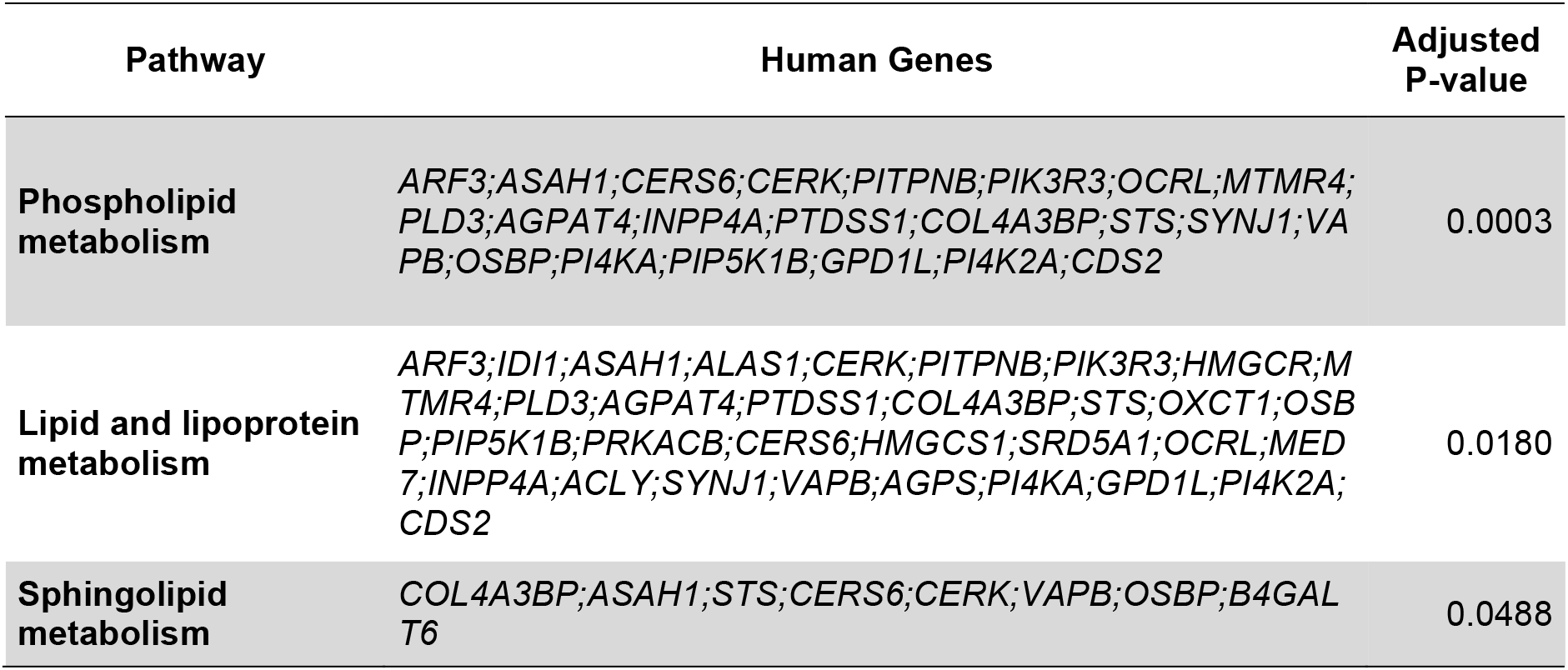
Genes involved in lipid metabolism. Genes down-regulated in the *substantia nigra pars compacta* of PD patients involved in lipid metabolism pathways.

### Glial expression of *spen* protects *Drosophila* from paraquat-induced neurotoxicity

We first investigated *spen* mRNA levels in the brain of adult *Drosophila* submitted to paraquat-induced toxicity. We could observe a significant up-regulation of *spen* expression by q-PCR in the brains of paraquat-treated flies (figure 1A). To get insights on the functions of Spen under such experimental conditions, flies heterozygotes for a *spen* loss-of-function mutation^15^ (*spen^**k07612**^/+* and *spen^03350^/+*) were submitted to paraquat intoxication. We found that the *spen* heterozygous mutant lines exhibit a higher sensitivity to paraquat as compared with control flies (figure 1B and supplemental figure 1). These results indicate that Spen may exert a protective role against paraquat-induced neurotoxicity. As previously reported in a study from Stein Aerts and collaborators^30^, we observed that *spen* expression, as revealed by PlacW inserts, is detectable both in neurons and in glial cells of *Drosophila* adult brain (supplemental figure 2). To determine whether the neuroprotective function of *spen* is linked to its expression in glial cells, we generated flies in which *spen* was selectively down-regulated in glial cells with a *UAS-spen*^*RNAi*^ and a pan-glial driver. We find that downregulation of *spen* in glial cells also leads to an exacerbated sensitivity to paraquat toxicity (figure 1C). Similarly, knockdown of *spen* specifically in adult glia, using the TARGET system^31^ increased *Drosophila* sensitivity to paraquat (figure 1D). Conversely, an over-expression of *spen* in glial cells had a protective effect (figure 1C). Collectively, these results show a protective role of Spen in the glial cells of *Drosophila* exposed to paraquat.

**Figure 1.**
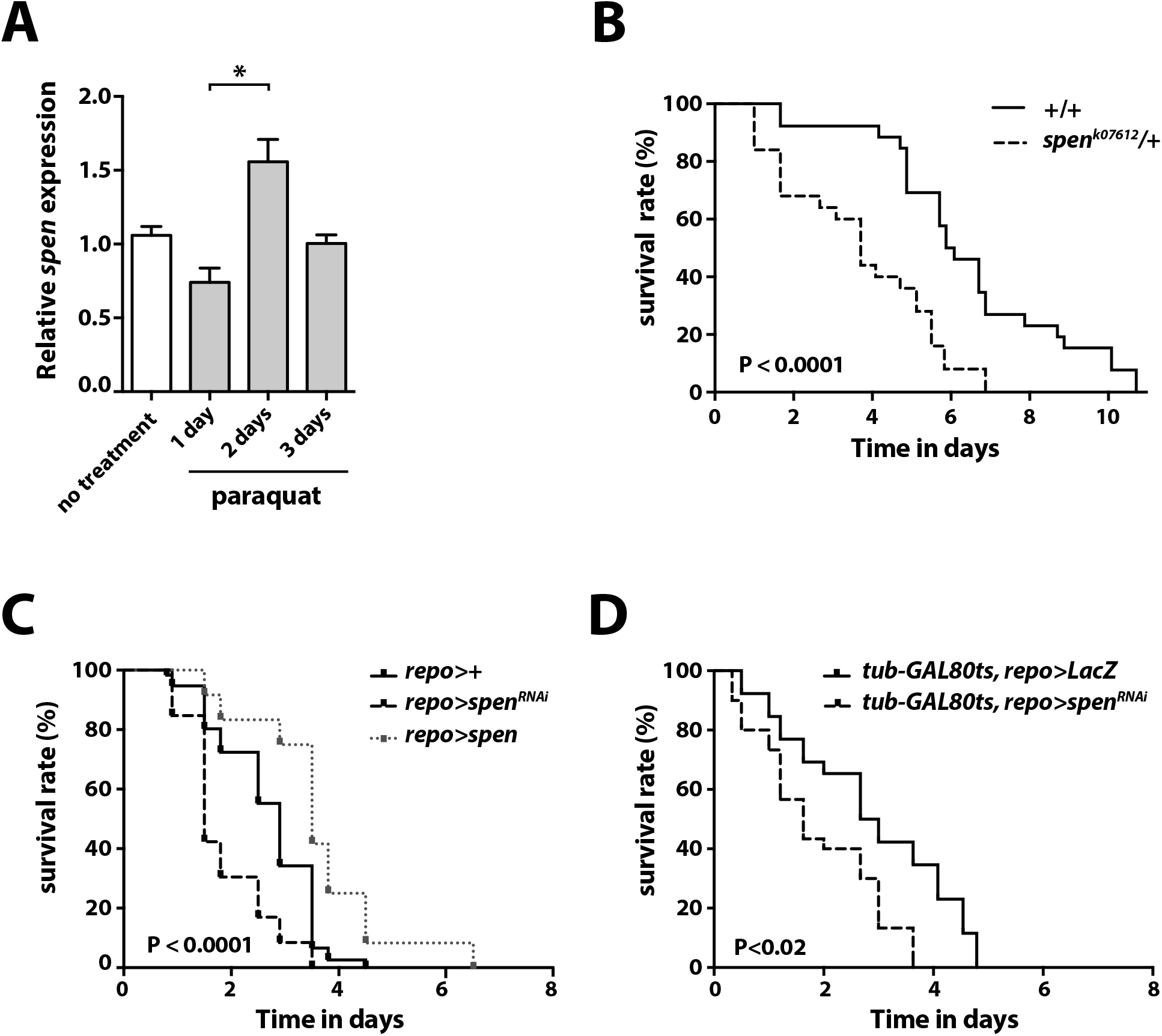
*spen* expression in glia is protective against paraquat-induced lethality. **(A)** Quantification of *spen* expression by q-PCR in heads of flies fed with a sucrose solution (no treatment) or a sucrose solution containing paraquat (10 mM) for 1,2 or 3 days (4 independent experiments). Results are expressed relative to no treatment condition. Non parametric Mann-Whitney test, * P<0.03. **(B)** Survival rate of *spen* loss of function heterozygous mutants (*spen^k07612^/+*) adult flies fed with paraquat (10 mM)*. spen*^*k07612*^ heterozygous flies are more sensitive to paraquat than wild-type flies (*w^1118^)*. Log-rank Mantel-Cox test, P<0.0001. 3 independent experiments, n=16-20 flies per genotype. **(C)** Survival rate of *spen*^*RNAi*^ flies fed with paraquat (10 mM). Flies expressing *spen*^*RNAi*^ in glial cells (*;UAS-spen^RNAi^/+; repo-GAL4/+*) are more sensitive to paraquat than control flies (*;;repo-GAL4/+*) and than flies over-expressing *spen* in glial cells (*;UAS-spen/+;repo-GAL4/+*). 3 independent experiments, n=20 flies per genotype. Log-rank Mantel-Cox test, P<0.0001. **(D)** Survival rate of flies expressing *spen*^*RNAi*^ fed with paraquat (10 mM). Expression of *spen*^*RNAi*^ was restricted to adult glia using the TARGET system^31^. Briefly, flies developed at 18°C to inhibit GAL4 activity and were switched to 29°C as adults to induce the expression of *spen^RNAi^*. Note that the kinetic of each experiment depends on the temperature and fly genetic backgrounds.

**Figure 2.**
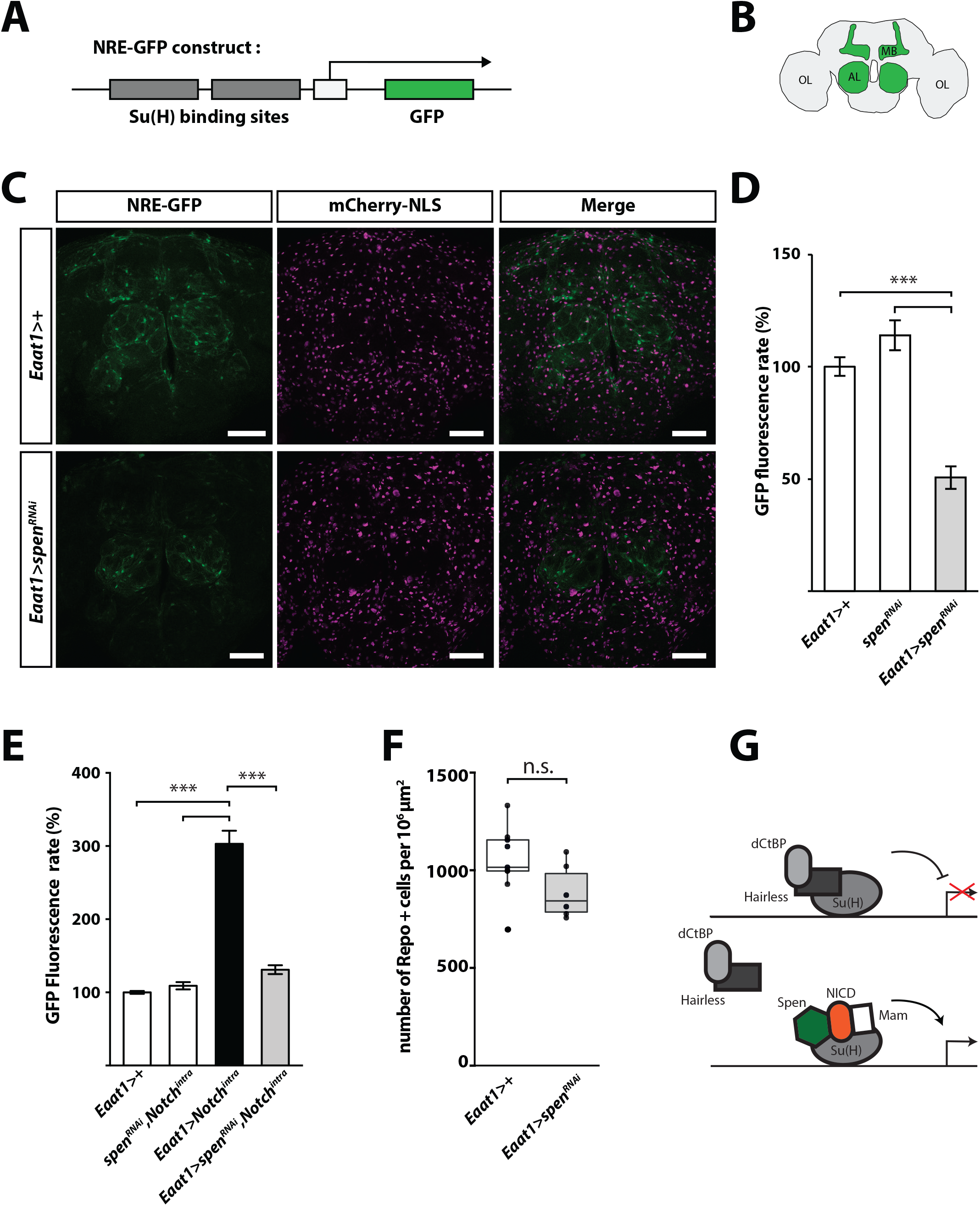
Spen regulates Notch signaling in *Drosophila* adult glial cells. **(A)** Notch activation was visualized in adult *Drosophila* brain using a Notch reporter (*NRE-GFP* construct) in which *EGFP* coding sequence (light gray) is under the control of a promoter containing multiple Su(H) binding sites (dark gray). **(B)** Simplified schematic of *Drosophila* adult brain, region known for Notch activity are coloured in green, namely the mushroom bodies (MB) and antennal lobes (AL). Optic lobes (OL). **(C)** NRE-GFP activation (right panel, green) in flies expressing *spen*^*RNAi*^ in glial cells was visualized by confocal microscopy in whole-mount brain. Glial cells expressing *Eaat1-GAL4*, are visualized with the nuclear marker *UAS-mCherry-NLS (*magenta). GFP-positive signal was detected in mushroom body (MB) and antennal (AL) lobes, in closed proximity with *Eaat1+* glial cells indicating that Notch activation occurred mainly in glia. NRE-GFP signal is reduced in Eaat1>*spen*^*RNAi*^ (;*Eaat1-GAL4/UAS-spen^RNAi^; UAS-mCherry-NLS/+*) compared to controls. Scale bar: 50μm. **(D)** Quantification of NRE-GFP fluorescence between control flies with driver alone *Eaat1>+ (;Eaat1-GAL4/+;NRE-GFP/+)* or UAS alone *spen^RNAi^ (UAS-spen^RNAi^/+; NRE-GFP/+*), and flies expressing *spen*^*RNAi*^ *Eaat1>spen*^*RNAi*^ (*;Eaat1-GAL4/UAS-spen^RNAi^; NRE-GFP/+)*. 3 independent experiments, n=18 flies per genotype. **(E)** NRE-GFP fluorescence was quantified in control driver alone *Eaat1>+* (*;Eaat1-GAL4/+; NRE-GFP/+)*, or UAS alone (;*UAS-spen^RNAi^, UAS-Notch^intra^;NRE-GFP/)* and compared with with Notch intracellular domain over-expression (*;Eaat1-GAL4/UAS-Notch^intra^; NRE-GFP/+)* alone or together with *spen*^*RNAi*^ (*;Eaat1-GAL4/ UAS-spen^RNAi^, UAS-Notch^intra^; NRE-GFP/+).* 3 independent experiments, n=18 flies per genotype. Non-parametric Kruskal-Wallis test, *** P<0.001 (in C and in D). **(F)** Average number of Repo positive glial cells per brain area in control *Eaat1>+* (*;Eaat1-GAL4/+;*) or *spen*^*RNAi*^ flies under the control of *Eaat1-GAL4* driver (*;Eaat1-GAL4/ UAS-spen^RNAi^;*) a minimum of 12 slices from 4 different brains were analyzed. **(G)** A simplified scheme of the Notch activator complex in adult glial cells would include Notch-intracellular domain (NICD), Suppressor of hairless (Su(H)), mastermind (Mam) and Spen. The Notch repressor complex includes Su(H), Hairless and dCtBP (*Drosophila* C-terminal Binding Protein). Adapted from ^51^.

### Spen regulates Notch pathway in adult *Drosophila* glial cells

Previous works showed that a crucial function of Spen and its human homolog, SPEN, is to regulate the Notch pathway, either positively or negatively depending on the molecular context and cell type considered. To determine if and how, Spen may regulate the Notch pathway in adult *Drosophila* glial cells, we performed *spen* knockdown using the Eaat1 (excitatory amino acid transporter 1) driver, which expression is mainly localized in astrocytes-like glial cells^30,32,33^. Notch pathway activation was monitored with a reporter transgene based on a minimal promoter containing Notch Responsive Elements, Su(H)-DNA binding sites, upstream of the *EGFP* coding sequence^34^ (*NRE-GFP* construct figure 2A). As previously reported, in control condition, we observed a basal expression of *NRE-GFP* and thus a basal activation of the Notch pathway, in adult glial cells, in particular in the antennal lobe^35^ (figure 2B, 2C and supplemental figure 3). Interestingly however, such a basal activation in glial cells was almost undetectable in flies expressing *spen*^*RNAi*^ in glia (figure 2C and 2D). This data indicates that Spen is required for basal Notch activation in glial cells of adult *Drosophila* brain. To further investigate the role of Spen in glial Notch signalling, we performed an epistasis experiment combining expression of the Notch intracellular domain, Notch^intra^, and *spen*^*RNAi*^ in Eaat1+ cells. As expected, Notch pathway activation by Notch^intra^ resulted in a strong induction of the of *NRE-GFP* reporter expression, and *spen* knockdown was sufficient to significantly reduce such an induction, indicating that Spen is necessary to maintain Su(H) dependent Notch signalling in adult glia (figure 2D, 2E). This positive regulation of Notch was not associated with a modification of glial cell survival rate as assessed by Repo staining quantification in control and *spen* knockdown condition (figure 2F). Collectively, these results show that Spen acts as positive regulator of Notch signalling in adult glial cells (figure 2G).

**Figure 3.**
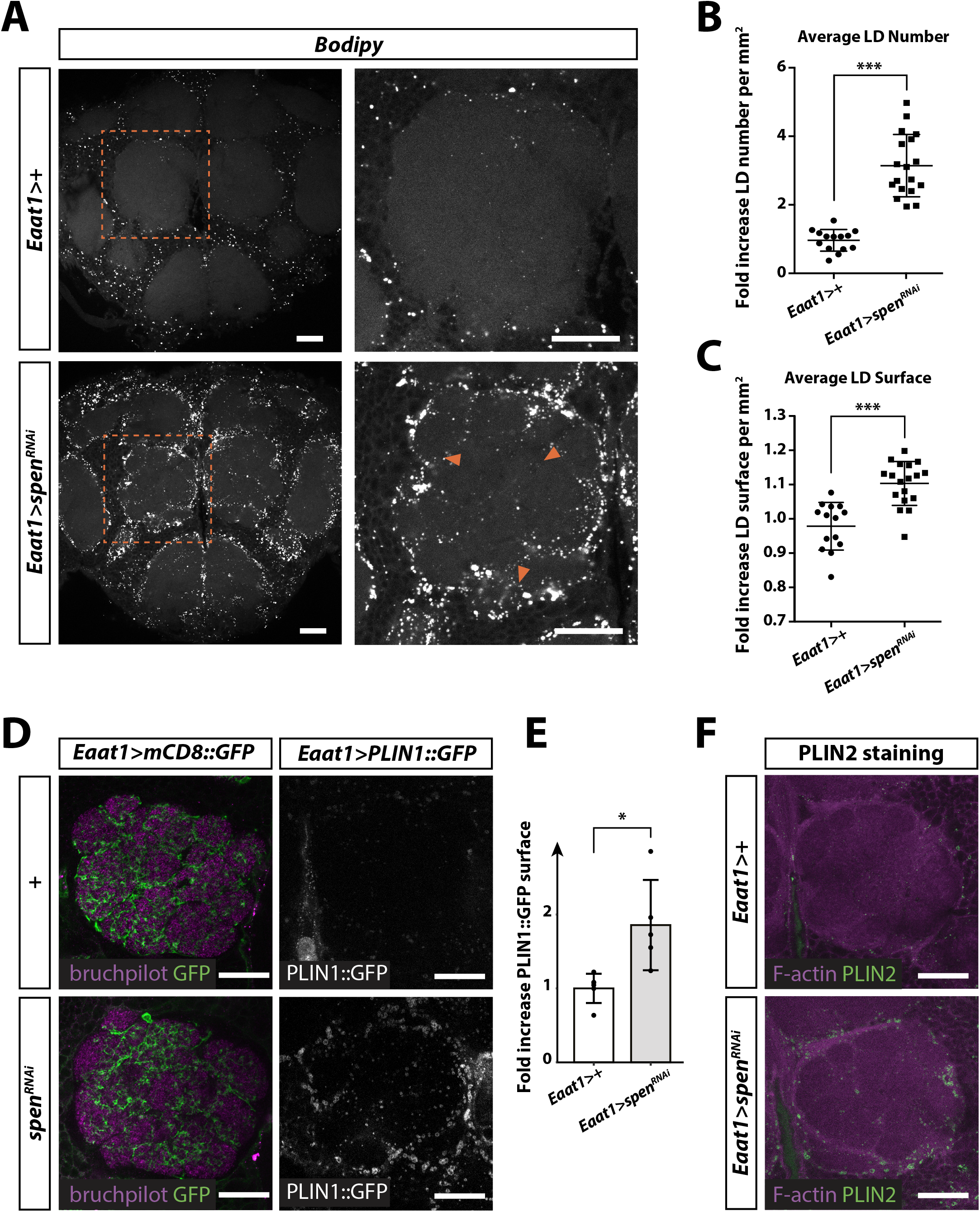
Spen regulates lipid droplets number, size and localization in glial cells. **(A)** Lipid droplets labelled with BODIPY 493/503 (white dots) in whole-mount brain of control flies *Eaat1>+ (;Eaat1-GAL4/+;*) or flies expressing *spen*^*RNAi*^ in glial cells *Eaat1>spen*^*RNAi*^ (*;Eaat1-GAL4/UAS-spen^RNAi^;*). As shown by the orange arrowheads in the close-up, lipid droplets are accumulating in the neuropil area of the antennal lobe in *spen* knockdown. Scale bar: 25μm. **(B-C)** Quantification of BODIPY 493/503 labelled lipid droplets. Data are presented as the fold change in lipid droplets number (B) and lipid droplet surface (C) in *Eaat1>spen*^*RNAi*^ relative to control flies *Eaat1>+* (;*Eaat1-GAL4/+;*). n=14–18 brains per condition. Droplets were quantified using an automated ImageJ plugin described in the material and methods section. Unpaired student T test, *** P<0.0001 **(D)** Lipid droplets surface were labelled with PLIN2 antibody staining (green) in whole mount brain of control flies *Eaat1>+* (; *Eaat1-GAL4/+;*) and *spen*^*RNAi*^ (;*Eaat1-GAL4/UAS-spen^RNAi^;*). Brains are counterstained with phalloidin-rhodamin (magenta). **(E)** Astrocyte-like glial processes are visible in the antennal lobe in flies expressing the membrane marker mCD8::GFP under the control of *Eaat1-GAL4* driver (left panel, *;Eaat1-GAL4/UAS-mCD8::GFP;).* Genetically encoded reporter of lipid droplet *UAS-PLIN1::GFP* (PLIN1::GFP) is labelling lipid droplet surface (right panel). As observed with the BODIPY staining in A, *spen* knockdown induces a lipid droplet accumulation in the antennal lobe where *Eaat1* positive glial cells extend their cellular processes. Scale bar: 25μm. **(F)** Quantification of PLIN1::GFP area fold change in *Eaat1>spen*^*RNAi*^ relative to control flies *Eaat1>+* (;*Eaat1-GAL4/+;*). n=5 brains per condition. Unpaired student T test, * P<0.05. Scale bar: 25μm.

### Spen regulates the number, size and localization of lipid droplets in glial cells

In *Drosophila* glial cells, lipid droplets were shown to accumulate under conditions of oxidative stress^5,6^. Interestingly, Spen has been shown to control lipid metabolism in the fat body of *Drosophila* larvae^10^. We thus investigated the impact of Spen on lipid droplets expansion and accumulation in *Drosophila* glial cells. To this aim, we developed an ImageJ^36^ macro to quantify lipid droplets in CNS cells of adult *Drosophila*. Briefly, quantification was based on an automated detection of fluorescent BODIPY positive particles that can be distinguished from background non-specific fluorescence by successive iterations. This new method allows not only discriminating lipid droplets from fluorescent foci doublets identified on successive confocal stacks but also assessing the density, size or circularity of lipid droplets (see the detailed description in the material and methods section). As shown in figure 3A and quantified in the antennal lobe neuropil (figure 3B and 3C), *spen* knockdown in glial cells induces a significant increase of lipid droplet number and size. These results were confirmed by the staining of whole-mount brains with an antibody directed against PLIN2^37,38^, a protein that is anchored in the phospholipid monolayer surrounding lipid droplets (figure 3D). In addition, we use the *UAS-PLIN1::GFP*, as a genetically encoded lipid droplet reporter^37,38^. Specific expression of PLIN1::GFP under the control of the *Eaat1-GAL4* driver further demonstrated a glial specific accumulation of lipid droplets in *spen* knockdown adult brain (figure 3E and 3F). Interestingly, Notch pathway inhibition by knocking down Su(H) specifically in glia did not affect lipid droplet number and size (supplemental figure 4). This suggests that lipid droplet accumulation in *spen* knockdown is not due to the inhibition of Notch signalling.

## DISCUSSION

Lipid dysregulation including lipid droplets dyshomeostasis have been reported to be part of glial cells’ stress response during neurodegeneration processes^5,7–9^ and in the context of PD^39^. Here, we found that *SPEN/SHARP* belongs to the set of astrocyte-expressed genes which mRNA levels are significantly different in the *substantia nigra* of PD patients as compared to controls. Interestingly, the list of differentially expressed genes was also enriched in genes involved in lipid metabolism. We showed that *spen* is up-regulated in the brain of paraquat treated flies, a model of PD in *Drosophila.* Moreover, *spen* mutant flies or flies expressing *spen*^*RNAi*^ in glia exhibited an increased susceptibility to paraquat. We also show that Spen is a positive regulator of Notch signalling in glial cells. Finally, we found that glial specific loss of *spen* led to the accumulation of lipid droplets in glial cells, which supports the central role of glial cells in the control of brain lipid metabolism.

*SPEN*, the human ortholog of *spen*, belongs to the list of significantly up-regulated astrocyte-expressed genes in PD patients. This is of particular interest in view of the previously reported role of Spen as a regulator of lipid storage in the adipocyte-like cells of *Drosophila*. Indeed, Spen was identified in two independent screens for its ability to modulate fat content in *Drosophila* larvae and adult fat body^40,41^. Moreover, in a more recent study, *spen* manipulation in adipocyte-like cells was correlated with the altered expression of key metabolic enzymes pointing to a role of Spen in energy catabolism^10^. Here, we found that, in a similar manner, the silencing of *spen* specifically in glial cells was responsible for an alteration of lipid metabolism, which translated into an increase of lipid droplet number and size in *Drosophila* glial cells. These results establish an important role of Spen in the control of lipid metabolism not only in adipocytes but also in adult *Drosophila* glial cells.

We also found that Spen is a positive regulator of Notch signaling in adult *Drosophila* glial cells. A Spen-dependent activation of the Notch pathway has also been observed in intestinal stem cells of adult flies^42^, further supporting that Spen can act as a positive regulator of Notch in adult tissues. This also raises the possibility that in *spen* mutant a reduced Notch signaling leads to the accumulation of lipid droplets although we could not modify lipid droplets size and number by knocking down Su(H) in glial cells. Rather, Spen may act directly at the level of mRNA stability or splicing to promote the expression of lipid metabolism genes independently of Notch^10^. Finally, the accumulation of lipid droplets may be due to an unleashed oxidative stress in *spen* mutant, as previously observed in flies with defective mitochondrial respiration or submitted to hypoxic treatments^5,6,8^. We may favor the latter hypothesis in light of our results showing that *spen* mutant flies exhibit a reduced resistance to paraquat-induced oxidative stress.

We also show that assessing the role of glia-expressed Spen in the lipid metabolism is relevant in the context of PD pathophysiology. Indeed, using a data mining approach, we found that genes previously shown to exhibit an altered expression profile in the *substantia nigra* of PD are not only enriched in astrocyte-expressed genes such as *SPEN/SHARP* but also include a significant number of genes annotated with the GO terms “phospholipid metabolism”, “lipid and lipoprotein metabolism” or “sphingolipid metabolism” (Table 1). These results pointing to a pathophysiological role of lipid metabolism in PD are in accordance with a previous data mining work based on the meta-analysis of GWAS (genome-wide association studies) studies^43^. Importantly, we found that several lipid-related genes, which are differentially expressed in the *substantia nigra* of PD patients, are likely involved in the regulation of lipid droplets formation and fate. This is notably the case for *AGPAT4* (*1-acylglycerol-3-phosphate O-acyltransferase 4*), which is involved in the synthesis of precursors of triglycerides, the major lipid component of lipid droplet. This holds true also for *SCD5* (*stearoyl-CoA desaturase 5*) a recently identified new target for PD treatment^44^, which catalyzes free fatty acid desaturation and play an important role in the early steps of lipid droplet formation. Finally, *Arf79F* and *schlank*, the *Drosophila* orthologs of two lipid-related genes found to be down regulated in the *substantia nigra* of PD patients were experimentally shown to control the homeostasis of lipid droplets^45,46^.

In conclusion, our data support a central role of *spen* expression in glial cells in the control of lipid metabolism and resistance to paraquat toxicity. Further investigation on the role of Spen will contribute to the understanding of the consequences of lipid dyshomeostasis in neurodegeneration.

## MATERIALS AND METHODS

### Fly strains and genetic

Flies were maintained on standard yeast medium at 25°C unless otherwise noted. Flies bearing the following mutations and transgenes were obtained from the Bloomington Drosophila Stock Center (Indiana University, USA): *w^1118^*, *repo-GAL4 (BL7415)*, *UAS-mCherry-NLS* (BL38424), *tubulin-GAL80*^*ts*^ (BL7019), *UAS-mCD8::GFP* (BL32186), *NRE-EGFP 1* (referred as NRE-GFP, BL30728), UAS-Notch^*intra*^ (BL52008), *Eaat1-GAL4*^32^ (BL8849). The *spen*^*k07612*^ and *spen*^*03350*^ P element insertion (DGRC, #102574 and BL11295) was previously characterized as homozygous lethal *spen* loss of function mutant^15^; they were used here as heterozygotes. *UAS-plin1::GFP*^37^ from R.P. Kuhnlein (University of Graz, Austria). An EP line, *spen-GS2279*, available at the Kyoto DGRC Stock Center was used to over-express *spen,* and is referred to as *UAS-spen*. The *UAS-spen*^*RNAi*^ was a gift from K.M. Cadigan^47^ and was previously characterized in *Drosophila* retina development^12^. *spen* mutants and transgenic flies were outcrossed to a *w*^*1118*^ control stock.

### RNA extraction and q-PCR

Total mRNA was isolated from 25 to 35 *Drosophila* heads following the RNeasy mini kit extraction protocol from Qiagen and reverse transcribed with oligo(dT)15 and the ImProm-II Reverse Transcription System (Promega) according to manufacturers’ instructions. Quantitative PCR reactions were carried out on a StepOnePlus (Applied Biosystems) using FastStart Universal SYBER Green Master (Roche Applied Science). Primers set efficiency (E) was assessed using serial dilutions of cDNA preparations. Standard curves were used to determine the relationship between mRNA abundance and PCR cycle numbers (Ct) and to calculate relative quantity (Qr = E−Ct)^48^. Values were normalized to Rp49 mRNA amount. Data from three independent experiments are presented as means ± standard deviation. q-PCR reactions were performed using the following primers for *spen*: 5’-TTCGTTGTGGGATAGCAGCA-3’; 5’-CGTTCGAAGCTGTTTGTCG-3’ and for *Rp49* (5’-ATCGTGAAGAAGCGCACCAAG-3’;5’-ACCAGGAACTTCTTGAATCCG-3’

### Immunostainings

*Drosophila* heads were cut off and kept in a drop of fresh Hemolymph Like 3 dissection buffer^49^ (HL3) supplemented with D-glucose (120 mM). Dissected brains were collected in a dish containing 1% PFA diluted in HL3 medium and fixed overnight at 4°C. Fixed brains were washed 3 times during 30 minutes in PBS containing Triton X-100 (0.5%), BSA (5 mg/ml), incubated 1h in PBS containing Triton X-100 (0.5%), BSA (5 mg/ml) and Normal Goat Serum (NGS, 4%) for blocking and then overnight with indicated primary antibodies: mouse anti-Repo 1/400 (DSHB, 8D12), rabbit anti-GFP 1/400 (Invitrogen, A6455) or rabbit anti-PLIN2^37^ (1/1000, Gift from R.P. Kuhnlein) diluted in blocking solution. After 3 washes, samples were incubated overnight with the appropriate secondary antibody: anti-mouse Alexa Fluor 647 1/400 (Invitrogen, A31571) or anti-rabbit Alexa Fluor 488 1/400 (Invitrogen A11008) diluted in blocking solution. Samples were washed 3 times and then mounted in Vectashield mounting medium (AbCys) on a bridge slide to prevent tissue flattening. Samples were stored at −20°C until visualization.

For NRE-GFP experiments, *Drosophila* heads were cut off and kept in a drop of fresh PBS. The proboscis was removed and cuticle opened in order to get access to the brain. After careful dissection brains were then transferred to 4% PFA containing PBS for 20 minutes at room temperature. After fixation, tissues were washed in PBS and mounted directly in Vectashield medium (AbCys). For these experiments wild type brains expressing no GFP were also dissected to evaluate the background fluorescence of the tissue.

### Image processing

Images were acquired at the LYMIC-PLATIM – Imaging and Microscopy Core Facility of SFR Biosciences (UMS3444, ENS de Lyon, France). For PLIN1::GFP, Repo and BODIPY staining, images were acquired on a Leica LSM800 confocal microscope and analyzed with the ImageJ^36^ software (see section below). For NRE-GFP, images were acquired on a Leica epifluorescence microscope. In each experiment, the mean GFP fluorescence of control brain (;*Eaat1-GAL4/+;*) was used to normalize the results. The NRE reporter is a synthetic construct with three copies of the SPS site (for Su(H) Paired Sites, binding sites for the Notch activity-dependent transcription factor, Su(H)) that is taken from the E(spl) regulatory region^34^.

### Automated Image analysis

We developed an ImageJ macro (https://gitbio.ens-lyon.fr/dcluet/Lipid_Droplets) to identify fluorescent particles on confocal stacks using *Drosophila* brains stained with the lipid droplet dye BODIPY 493/503 (Molecular Probes, D-3922) or labelled with the glial specific marker Repo (immunlocalization of glial nuclei). The program requires ImageJ^36^ v1.49g or higher and is based on an iterative detection of the brightest particles followed by the removal of “doublets” of the same particle over the stack. Briefly, the program first identifies the tissue all along the stack. The particles will then be sought within the tissue all along the stack. The signal is first intensified using the Gaussian blur function and the maximum entropy treatment. The “Max-Entropy” threshold method^50^ then allows the detection of the particles of interest. The detected particles are stored in a transient matrix, and removed from the image. Thus, the next iteration is able to detect less bright particles. Finally, the program removes all doublets of the same particle along the z-dimension of the stack (keeping the largest candidate as the best) to have an optimal counting of the labelled particles. Finally, multiple parameters can be calculated, such as particule density, size or circularity.

### Paraquat-induced PD model

Paraquat medium was prepared shortly before use as follows: paraquat (Sigma, 36541) was added to 0.8% low melting agarose (Sigma, A9414), 10% sucrose (Sigma, S0389), in PBS solution. 3-days old flies were starved during 4 hours on 0.8% agarose medium and then fed with 10 mM paraquat containing medium for 5-7 days in survival experiments or indicated times for RNA extractions.

At least three independent experiments were performed for each shown data set with n≥20 flies per condition for each experiment.

### Lipid droplet staining

Heads of 6 days old flies were dissected in a drop of fresh HL3 dissection buffer supplemented with D-glucose (120 mM). Dissected brains were collected in a dish containing 1% PFA diluted in HL3 medium and then fixed overnight at 4°C. Brains were washed 3 times during 30 minutes with PBS supplemented with Triton X-100 (0.1%) and then incubated overnight with BODIPY 493/503 (Molecular Probes, D-3922) 15 μg/mL diluted in PBS, Triton X-100 (0.1%) at 4°C. Subsequently, brains were washed 3 additional times with PBS, Triton X-100 (0.1%) and mounted in Vectashield medium (AbCys) on a bridge slide and stored at −20°C until imaging.

## ACKNOWLEDGMENTS

This work was supported by the French National Research Agency award ANR-12-BSV1-0019 to BM, « Projets développement technologique » SFR Biosciences to ND and BM. ARTHRO-TOOLS and PLATIM facilities. VG PhD thesis was supported by grants from Fondation Servier, ENS fond recherche and a FRM fellowship. MQ PhD thesis was supported by fellowships from the French Ministry of Research and education and France Parkinson.

## AUTHOR CONTRIBUTIONS

V. Girard, V Goubard, M Querenet, E. Dufourd, and Nathalie D performed the experimental work using *Drosophila*. L. Seugnet designed and performed the Notch experimental part of the article with the help of V. Goubard. L. Pays and S. Nataf did the data mining study. D. Cluet set up the Image J Macro. N. Davoust designed the experiments and wrote the manuscript. V. Girard, S. Nataf, L. Seugnet and B. Mollereau contributed to the writing of the manuscript as well.

