## Supplemental figures 1-4 for "Split-ends is modulating lipid droplet content in adult *Drosophila* glial cells and is protective against paraquat toxicity"

### Supplemental Files

### Girard et al, Supplemental Figure 1

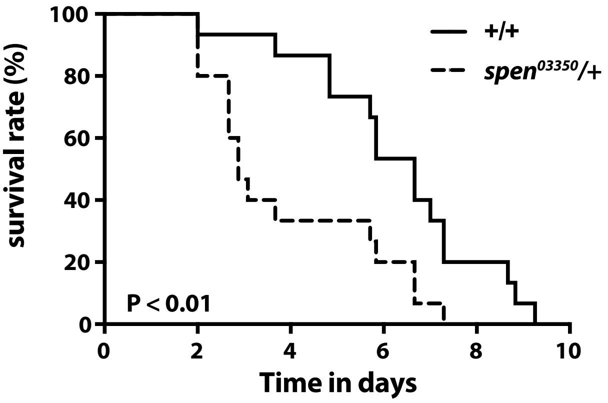


**Figure S1.** ***spen*** **heterozygous mutant flies are more sensitive to paraquat-induced lethality.** Survival rate of *spen* loss of function heterozygous mutants (*P{lacW}spen[03350]/+*) adult flies fed with paraquat (10 mM). *spen*$^{03350}$ heterozygous flies are more sensitive to paraquat than wild-type flies (*w1118*). Log-rank Mantel-Cox test, P<0.0001.

### Girard et al, Supplemental Figure 2

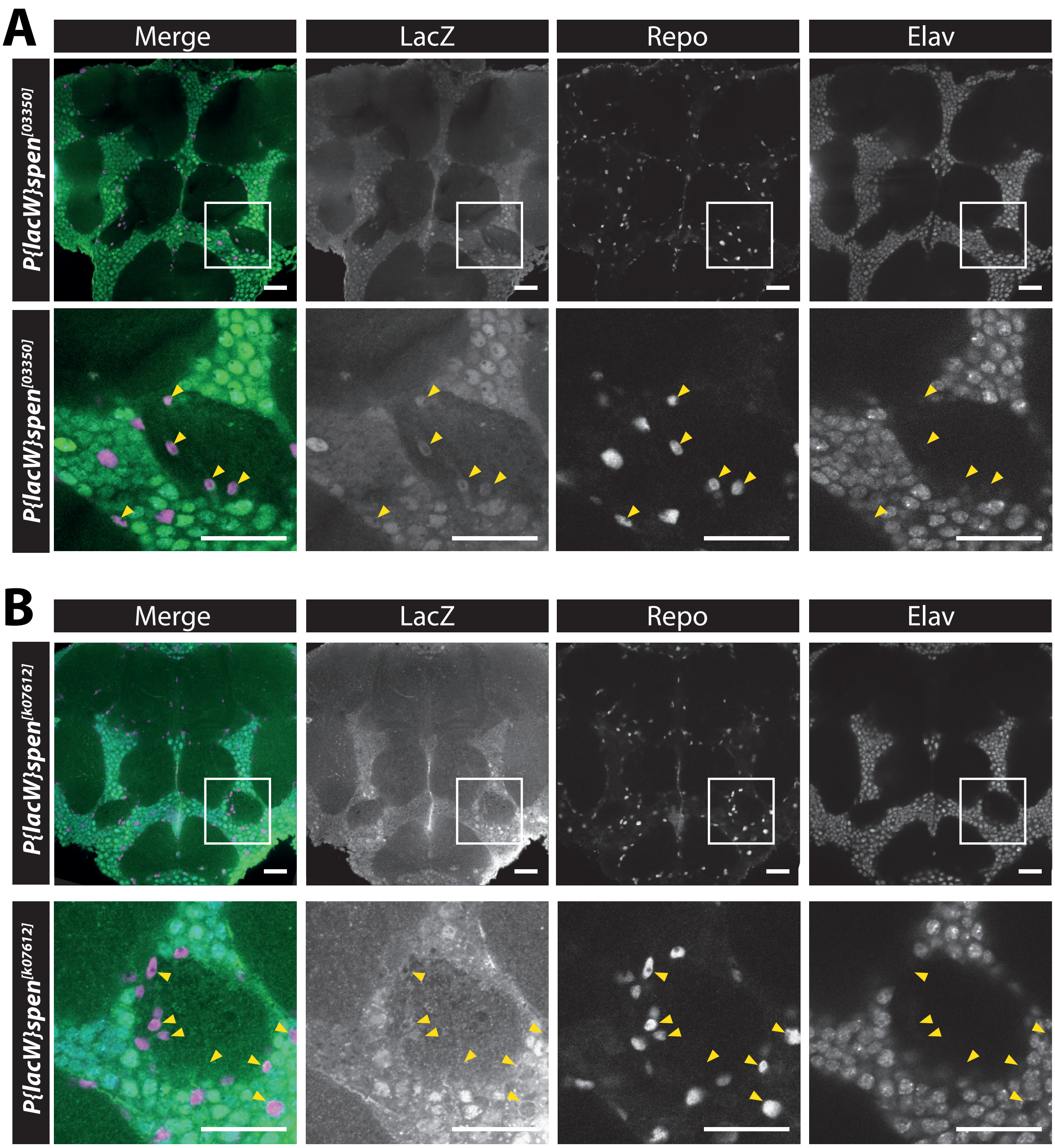


**Figure S2.** ***spen*** **is expressed in both glia and neurons in** ***Drosophila*** **adult brain.**
(A) *spen* expression domain in adult brain was assessed by immunostaining of LacZ (green) reporter gene in whole mount brain of 2 enhancer trap lines: *P{lacW}spen[03350]*, **(A)** and *P{lacW}spen[k07612],* **(B)**. Glial cell nuclei are visualized using Repo staining (magenta) and neuron nuclei using Elav staining (cyan). Expression of *spen* in glial cells is indicated with yellow arrowheads. Scale bar: 25µm.

### Girard et al, Supplemental Figure 3

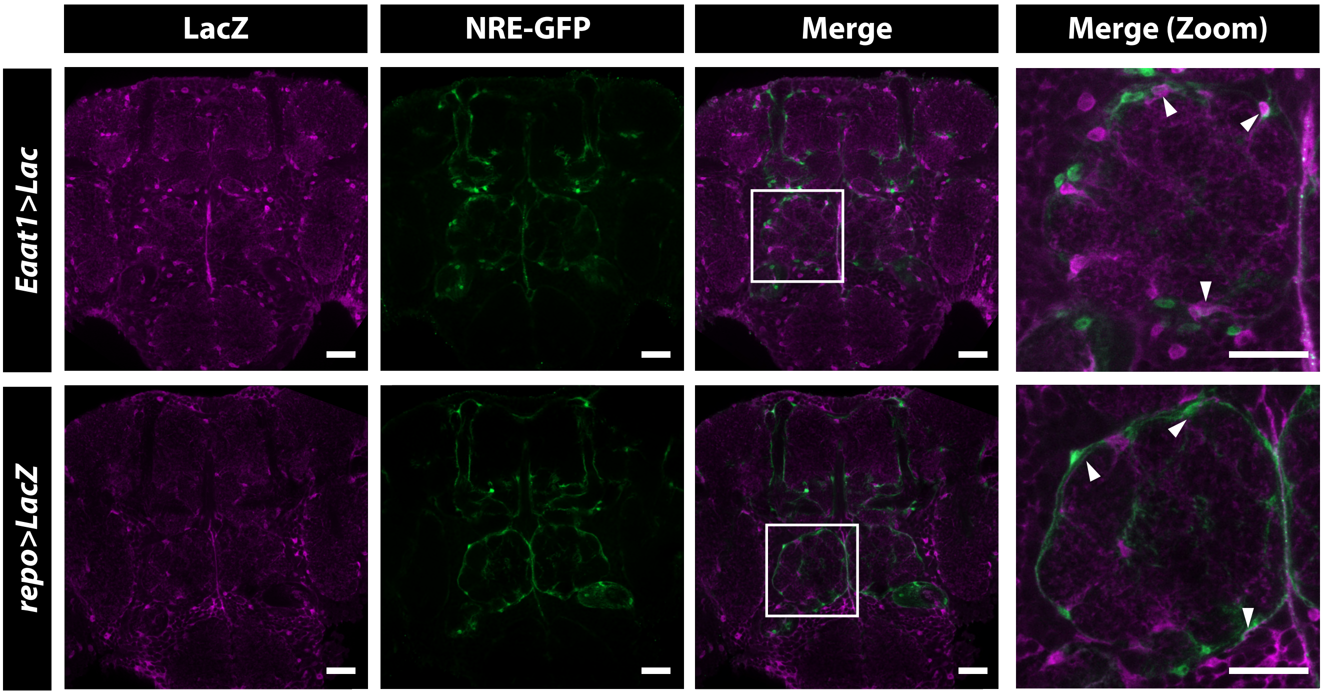


**Figure S3. Notch signalling is visible in adult glial cells expressing Eaat1-Gal4.**
Confocal microscopy of whole-mount brain of flies carrying Notch activation reporter NRE-GFP (green) expressing LacZ (magenta) under the control of glial driver *Eaat1-Gal4* and *repo-Gal4*. NRE-GFP and LacZ co-localisation in glial cells Eaat1 or repo positive is indicated with white arrowheads. Scale bar: 25µm.

### Girard et al, Supplemental Figure 4

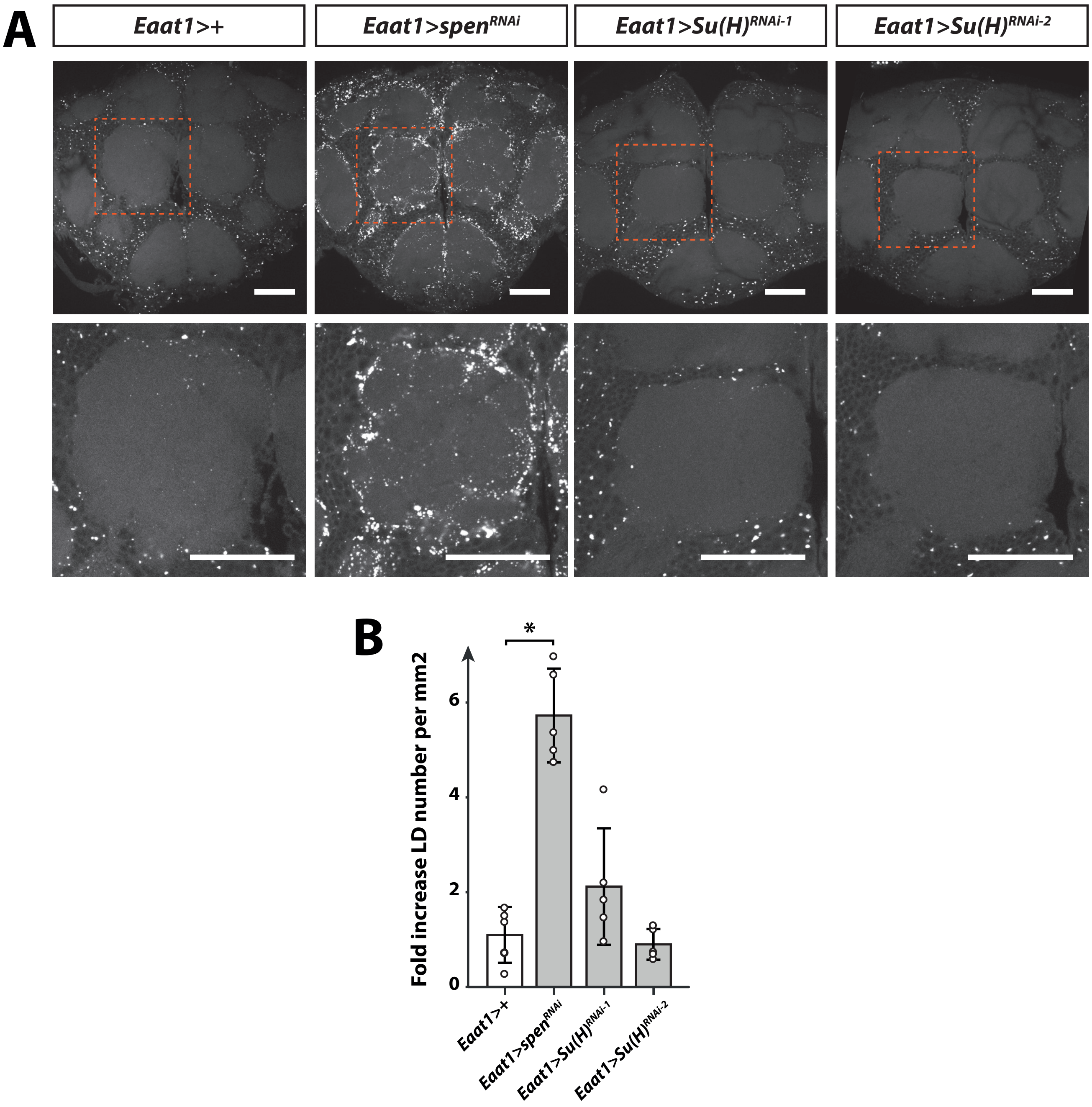


**Figure S4. *Su(H)* knockdown do not modify glial cell lipid droplet in adult *Drosophila* brain. (A)** Lipid droplets labelled with BODIPY 493/503 (white dots) in whole-mount brain of control flies *Eaat1>+* (*;Eaat1-GAL4/+;*) and flies expressing $spen^{RNAi}$ in glial cells $Eaat1{>}spen^{RNAi}$ ($;Eaat1\text{-}GAL4/UAS\text{-}spen^{RNAi;}$) was compared to flies expressing two different $Su(H)^{RNAi}$ (RNAi-1 : HMS05110 , RNAi-2: KK101550). As shown in the close-up, lipid droplets are accumulating in the neuropil area of the antennal lobe in *spen* knockdown but not in *Su(H)* knockdown. Scale bar: 25µm. **(B)** Quantification of BODIPY 493/503 labelled lipid droplets. Data are presented as the fold change in lipid droplets number (B) relative to control flies *Eaat1>+* (;Eaat1-GAL4/+;). n=5 brains per condition. Droplets were quantified using an automated ImageJ plugin described in the material and methods section. Non-parametric Kruskal-Wallis test, * P<0.05. Scale bar: 50µm.
